# Comparative analysis of two autophagy-enhancing small molecules (AUTEN-67 and -99) in a *Drosophila* model of Spinocerebellar ataxia type 1

**DOI:** 10.1101/2025.09.18.677017

**Authors:** Tímea Burján, Maryam Aslam, Fanni Keresztes, Tímea Sigmond, Viktor A. Billes, Norbert Bencsik, Katalin Schlett, Tibor Vellai, Tibor Kovács

## Abstract

Autophagy is a lysosome-mediated self-degradation process of eukaryotic cells which is critical for the elimination of cellular damage. Its capacity progressively declines with age, and this change can lead to the development of various neurodegenerative pathologies including Spinocerebellar ataxia type 1 (SCA1). SCA1 is mainly caused by mutations in the polyglutamine region of Ataxin 1 protein. In patients affected by the disease, Purkinje neurons of the cerebellum frequently undergo demise and eventually become lost. Here we tested whether two well-characterized autophagy-enhancing small molecules, AUTEN-67 and -99, which antagonize the autophagy complex Vps34 through blocking the myotubularin-related lipid phosphatase MTMR14/EDTP, have the capacity to ameliorate SCA1 symptoms. We found that in a *Drosophila* model of SCA1, only AUTEN-67 exerts positive effects including improvement of climbing ability and extending life span. Based on these results, we hypothesized that the two compounds influence autophagy in the brain in a neuron-specific manner. Indeed, according to data we obtained, AUTEN-67 and -99 exhibit shared and unique functional domains in the *Drosophila* brain. AUTENs enhanced autophagy in GABAergic and dopaminergic neurons. In addition, AUTEN-67 also affected autophagy in cholinergic neurons, while AUTEN-99 triggered the process in glutaminergic neurons and motoneurons. We also observed varying efficiencies between the two AUTENs among different subtypes of cultured hippocampal neurons of mice. These data suggest that the two compounds display neuron-specific differences in exerting autophagy-enhancing effects, and may lead to a better understanding of in which types of neurons autophagy could potentially be activated to treat SCA1 in human patients.

## Background

Defects in the autophagic process (as a consequence of inactivating mutations in *Atg*-related genes) can lead to the development of various age-associated diseases, such as sarcopenia, type II diabetes, cancer and certain neurodegenerative pathologies [1–3]. The capacity of autophagy declines with age in various, distantly related organisms [4–8] which leads to the progressive accumulation of cellular damage over time. Such damages interfere with cellular processes, thereby causing a functional decline (senescence) and, eventually, loss of the affected cell. When cell death occurs in large quantities, the affected tissue/organ also undergoes a functional deterioration, leading to the incidence of a fatal, age associated degenerative disease [9]. The operation of this cellular maintenance process is even more important in tissues where cells no longer divide and hence cannot substitute those being lost. For example, the nervous system largely consists of terminally differentiated neurons, the quantity of which strongly affects aging [10, 11]. Due to these reasons, autophagy has an important therapeutic potential in medicine. Thus, drug candidates with potent autophagy-inducing effects are now placed into the center of current pharmacological research. To date, several autophagy-inducing candidates were identified and characterized, many of which act upstream of the process. For example, rapamycin, which is frequently used as an immunosuppressant agent, antagonizes the mTORC1 complex (mechanistic kinase target of rapamycin complex 1). mTORC1 inhibits autophagy by repressing Atg1 kinase required for the activation of the process. mTORC1, however, influences many other cellular processes, such as translation, ribosome biogenesis, mitochondrial respiration, as well as lipid and nucleotide synthesis [12, 13], and this way the application of rapamycin can often lead to undesired side effects [14]. To isolate more effective autophagy enhancers is a current challenge in pharma industry.

Macroautophagy (hereafter referred to as autophagy) can be divided into five main stages: activation, phagophore nucleation (generation of the initial isolation membrane), autophagosome formation (a double bilayer membrane-bound structure that sequesters the cargo destined for degradation from the rest of the cytoplasm), fusion of the autophagosome with a lysosome (this compartment contains hydrolytic enzymes including proteases, lipases, nucleases and glycolases) to form an autolysosome, and, eventually, degradation of cytoplasmic materials in the autolysosome. The end-products (monomers and energy) of autophagic degradation can be used in the synthetic processes and for cellular functions. The stages are controlled by Atg (autophagy-related) proteins. Both mechanism and regulation of autophagy are highly conserved during evolution [15].

Phagophore nucleation relies on the availability of phosphatidylinositol 3-phosphate (PIP3), an important component of the isolation membrane (PIP3-containing membranes contribute to the formation of autophagic structures) [16]. PIP3 is generated by phosphorylating phosphatidylinositol (PI). The reaction is catalyzed by the Vps34/PI3K kinase complex [17]. PIP3 then binds to proteins containing a FYVE domain. The FYVE zinc finger domain is named after the four cysteine-rich proteins, Fab 1 (yeast ortholog of PIKfyve), YOTB (Y-box binding protein), Vac 1 (vesicle transport protein), and EEA1 (Early Endosome Antigen 1), in which it has been found [18]. Such a protein is Atg18 (the yeast ortholog of mammalian WIPI1/2) that recruits autophagy-related protein complexes (e.g., the Atg12-Atg5-Atg16 complex) to phagophore [19]. Mammalian myotubularin-related lipid phosphatase MTMR14/Jumpy and its *Drosophila* counterpart EDTP function to antagonize the Vps34/PI3K complex. These enzymes dephosphorylate PIP3 into PI, thereby protecting the cell from undergoing excessive autophagic self-degradation [20, 21]. The expression of MTMR14 and EDTP progressively increases in brain neurons during the adult life span in both humans and *Drosophila* [15]. As a consequence, age-related decline in the capacity of autophagy is more prevalent in this organ [8]. Indeed, inhibiting EDTP specifically in the nervous system promotes longevity and improves climbing ability in flies [8].

We previously identified and characterized two autophagy-enhancing small molecules, AUTEN-67 and -99, which interfere with MTMR14 and EDTP proteins. The two agents extend life span and enhance, especially at advanced ages, flying ability in *Drosophila*. AUTEN-67 provides a positive effect in a *Drosophila* model of Huntington’s disease (HD), whereas AUTEN-99 does the same in both HD and Parkinson’s disease (PD) models: the compounds increase moving ability, decrease protein aggregation, and inhibit neuronal demise in these systems [22–24]. Type I spinocerebellar ataxia (SCA1) is a fatal neurodegenerative pathology that is caused by the repetition of polyglutamine tracts within Ataxin1 protein. In humans, the disease manifests when the number of glutamine repetitions exceed 40 (40Q). The higher the number of repetitions is, the more severe the disease’s symptoms are [25, 26]. In this study we aimed to compare further the AUTEN molecules in the SCA1 neurodegenerative disease model, revealing additional details of the neuronal subtype-specific actions of the AUTEN compounds.

## Results

### AUTEN-67 activates autophagy in the nervous system of *Drosophila* SCA1 models

By using an *elav-Gal4* driver, we first expressed normal (30Q) and mutant (82Q) human ATXN1 proteins in *Drosophila* [27]. A frequently used autophagy marker is Atg8a that is a fly ortholog of human LC3B (light chain 3 B) [28]. We used an mCherry-Atg8a reporter construct that labels essentially all autophagic structures (from early phagophores to late autolysosomes) [29]. We found that in the 82Q mutant genetic background, AUTEN-99 significantly enhanced the amount of mCherry-Atg8a-poistive structures as compared with untreated control (82Q, DMSO) (Figure S1A and 1A’). Using a 2xFYVE-GFP marker, we could monitor the levels of PI3P, which is associated with the activity of the Vps34 complex. When EDTP is inhibited (by AUTEN treatments), we expected a significant increase in the number of 2xFYVE-GFP-positive structures [30]. Our results indicate that only AUTEN-67 significantly increased the quantity of 2xFYVE-GFP structures in 82Q ATXN1 mutants expressing *elav-Gal4* throughout the entire adult brain (Figure S1B and B’). The efficiency of acidic degradation was then examined by using an anti-ubiquitin-specific antibody labeling and a GFP-p62 reporter system (fluorescence microscopy). Ubiquitinated proteins and SQSTM1/p62 [sequestrome 1 in mammals and Ref(2)P in *Drosophila*] are considered substrates of autophagic breakdown, so their amounts are inversely related to autophagic activity [31]. The two AUTEN molecules showed a marked difference in modulating acidic breakdown in these systems. AUTEN-67, but not AUTEN-99, lowered the levels of p62/Ref(2)P-positive structures and ubiquitinated in brain samples of animals expressing 82Q (Figure 1A, A’, B and B’). We also performed a Western blot analysis to quantify p62/Ref(2)P and Atg8a protein levels in samples isolated from *Drosophila* heads [32]. In this assay, the amount of p62/Ref(2)P was elevated in 82Q mutant animals as compared with control (30Q animals). High levels of these proteins in 82Q animals were significantly reduced by treating with either of the two AUTEN molecules, but AUTEN-67 appeared to be more effective in this respect (Figure 1C and Figure S1C). Western blot analysis is suitable for distinguishing the two forms of Atg8a, cytoplasmic (soluble) Atg8a-I isoform and membrane-conjugated Atg8a-II isoform [33]. We found that both AUTEN-67 and -99 are each able to enhance the amount of Atg8a-II (Figure 1C). Taken together, both AUTEN molecules can influence autophagic activity in the brain of the *Drosophila* SCA1 model used, but AUTEN-67 is more effective in this function as compared with AUTEN-99.

**Figure 1.**
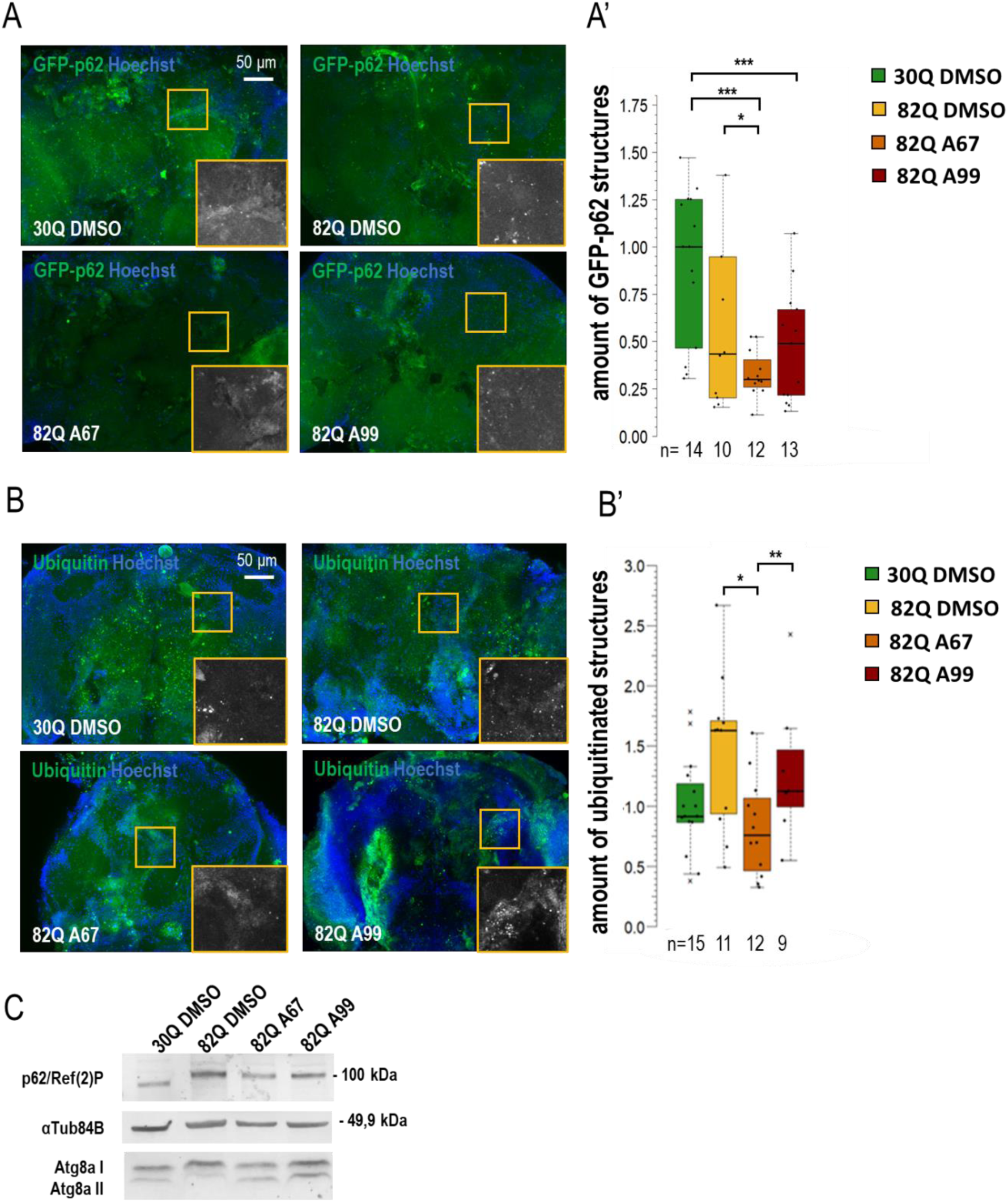
AUTEN-67 activates autophagy in a *Drosophila* model of SCA1. Determining autophagic activity in the brain of SCA1 model animals expressing wild-type (30Q) and mutant (82Q) human ATXN1 proteins. (**A**) GFP-p62/Ref(2)P reporter was used to quantify the level of autophagic degradation. 82Q ATXN1-expressing neurons were compared with 30Q ATXN1-expressing control neurons. Control was treated with DMSO (AUTEN molecules were dissolved in DMSO). Yellow squares indicate the enlarged areas. **(B-B’)** Determining protein aggregation by anti-ubiquitin labeling (green). Only AUTEN-67 (A67) could significantly lower the amount of ubiquitin-positive structures. Hoechst staining was used to visualize nuclei. **(C)** Western blot analysis on head samples obtained from animals at age of 21 days, and maintained at 29°C. Ref(2)P/p62 levels increase in 82Q animals as compared with control and 30Q samples. Both AUTEN molecules lower substrate levels but A67 does it with more effectively. The ratio of Atg8a-I and II isoforms in control vs. treated samples. A67 significantly changes the ratio. On the box-plot diagrams, the black line indicates the median value, the box shows the most representative 50%, while filed blacked circles display the value of individual samples. Stars represent P values, where *: p < 0,05, ***: p < 0,005. Detailed description of the statistical methods used can be found in the Materials and Methods chapter.

### AUTEN-67 improves movement and extends life span in SCA1 model flies

Similar to its human counterpart, a *Drosophila* model of SCA1 can be characterized by a decreased reaction ability (climbing) and a limited life span (Figure 2A and 2B, Figure S2A and B) [34]. The 30Q (control) and 82Q (SCA1 model) expressing flies were placed in thin tubes. After tapping, the animals start climbing up the sides of the glass from the bottom of the tube (negative geotaxis). This experiment allows the study of the animals’ response time (short-term) and the maintenance of the induced movement (long-term). Mutant animals had significantly worse climbing ability than the 30Q control, and the 82Q animals also had significantly shorter lifespans (Figure 2B). We found that the two AUTEN molecules improve climbing ability and promote longevity in 30Q flies (Figure 2A and B, Bi and Fig. S2). Contrary to these results, only AUTEN-67 was able to exert such positive effects in 82Q animals, while AUTEN-99 remained ineffective in this model (Figure 2A and B, Fig. S2).

**Figure 2:**
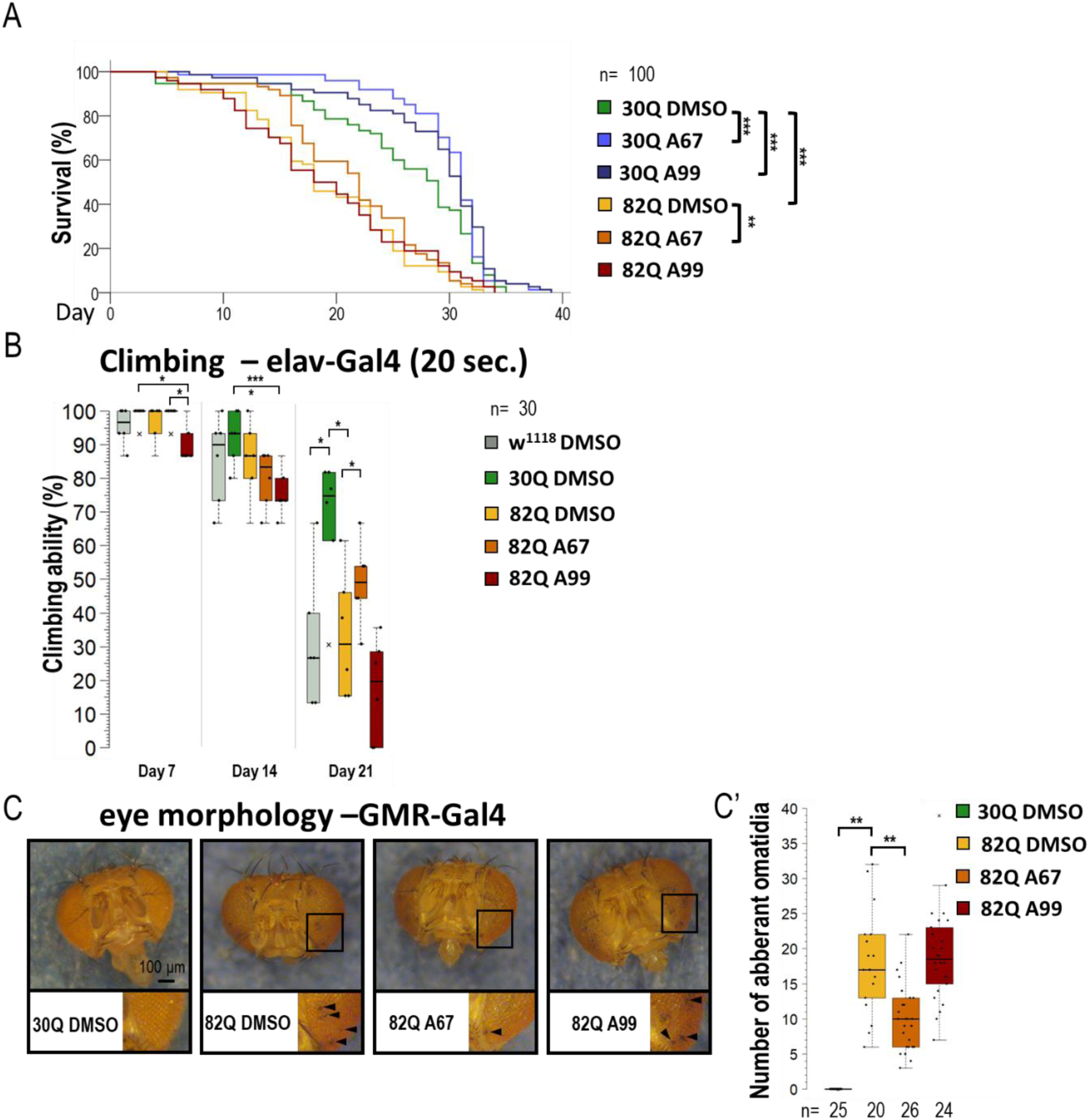
AUTEN-67 is more effective than AUTEN-99 in inhibiting the incidence of pathologies associated with SCA1 model. **(A)** The effect of AUTEN-67 (A67) and AUTEN-99 (A99) autophagy enhancing small molecules on life span in SCA1 models expressing wild-type (30Q) and mutant (82Q) ATXN1 proteins. In 30Q animals, both compounds significantly promoted longevity. However in 82Q animals, only A67 provided a life span advance. Control animals were treated with DMSO only (A67 and A99 were dissolved in DMSO). **(B)** Comparing climbing ability in control vs. SCA1 model animals. Only A67 could improve the ability of aged 82Q animals to move. **(C-C’)** Only A67 reduces the number of abnormal omatidia (indicated by black arrows) characterizing SCA1 models. On the box-plot diagrams, the black line shows the median value, the box indicates the most representative 50%, while filed black circles display the value of individual samples (when climbing was measured, the percentage of 10 animals). Stars indicate P values where *: p < 0.05, **: p<0.01, ***: p < 0005. Detailed description of the statistical methods used can be found in the Materials and Methods chapter.

SCA1 model (mutant 82Q ATXN1) animals are known to exhibit an aberrant compound eye morphology coupled with retina degradation [35, 36]. We indeed identified this characteristic eye phenotype in 82Q animals (Figure 2C and C’, Figure S2C and C’). For eye morphology studies, animals were treated under two conditions. First, larvae were treated on a medium containing AUTEN molecules (in nutrient), and after hatching, the ratio of aberrant ommatidia was counted (Figure S2 C and C’). Second, animals were treated with AUTENs for one week after hatching (Figure 2C and 2C’). In good accordance with data above, AUTEN-67, but not AUTEN-99, could lower the number of abnormal omatidia in the SCA1 model used (Figure 2C, C’ and Figure S2C, C’).

### AUTEN-67 activates autophagy in cholinergic neurons, whereas AUTEN-99 enhances the process in glutamatergic and motor neurons

Results obtained from the movement, life span and eye morphology assays suggest that the effects of the two AUTEN molecules on autophagy are different in distinct brain regions. This prompted us to select five specific *Gal4* drivers that are expressed in different types of neurons: *Appl-Gal4* (it is expressed in the whole brain except for the eye lobe) [37], *ple-Gal4* (it is expressed in dopaminergic neurons only [38], *Gad1-Gal4* (it is specific to GABAergic neurons) [39], *Chat-Gal4* (it is active in cholinergic neurons) [40], and *OK371-Gal4* (it is expressed in glutamatergic neurons and motoneurons) [41]) (Figure S3A). To monitor autophagic activity, we selected a UAS-GFP-mCherry-Atg8a (UGMA) reporter that is expressed in specific brain structures depending on the corresponding *Gal4* driver we used (this double reporter can distinguish early and late autophagic structures – phagophores and autophagosomes appear as yellow structures while autolysosomes are visible as red structures due to the fact that GFP is sensitive to the acidic milieu such asof the lumen of autolysosomales lumen). Using fluorescence microscopy, we assessed the amount of mCherry-positive autolysosomes only. We also used a UAS-GFP-2xFYVE (GFP-2xFYVE) reporter to monitor changes in PIP3 levels (Figure 3, 4 and Figure S3) [42]. To study how the two AUTEN molecules influence autophagy in specific neurons, a Western blot analysis was applied where the UGMA strain was stained with a GFP-specific antibody. In this experiment, acidic breakdown was quantified by monitoring the levels of GFP and GFP-mCherry proteins (Figure 3, 4 and Figure S3) [43].

**Figure 3:**
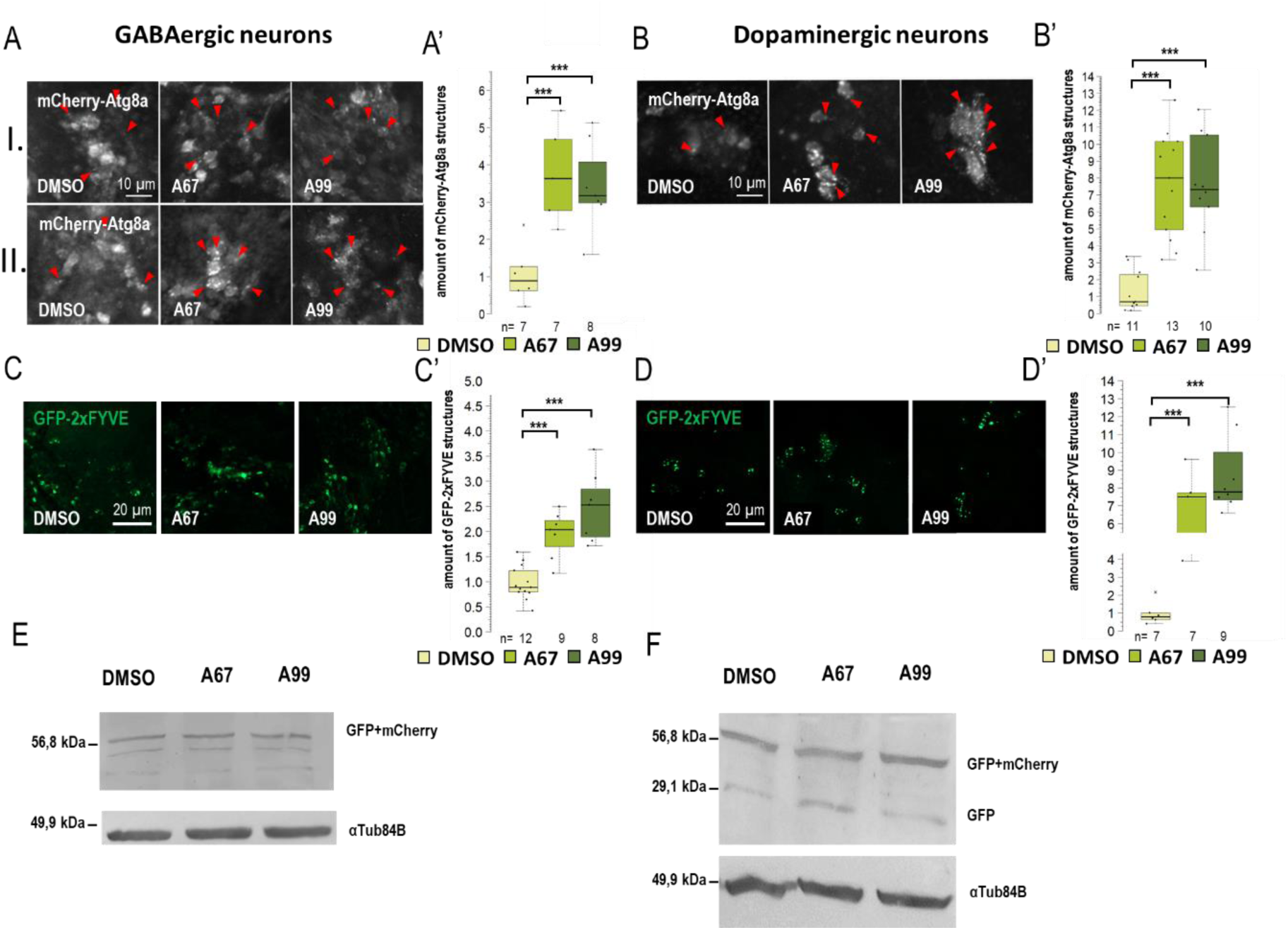
Both AUTEN-67 and -99 enhance autophagy in GABAergic and dopaminergic neurons. **(A-B)** mCherry**-**Atg8a reporter labeling autophagy vesicles was expressed in GABAergic and dopaminergic neurons by using specific Gal4-drivers. mCherry signals were detected by confocal microscopy. We examined the amount of autophagic vesicles in the brains of adult animals in two areas: (I) the mushroom body and (II) the neurons around the pharynx. Both AUTEN-67 (A67) and AUTEN-99 (A99) treatments increased the level of mCherry-Atg8a-positive structures. **(A)** In GABAergic neurons, cell groups were identified in the mushroom body (I.) and in the area under the pharynx (II.). Indicated areas are seen in Figure S3A, the analysis is shown in Figure S4A. **(C-D’).** PIP3 levels were determined by using a GFP-2xFYVE reporter. The reporter was expressed under the control of specific Gal4-drivers. A67 and A99 treatments increased the amount of PIP3-positive structures in GABAergic and dopaminergic neurons. **(E-F)** GFP-mCherry-Atg8a reporter was expressed in a neuron type-specific manner. On the Western blot, anti-GFP labeling distinguishes GFP-mCherry-Atg8a depredated forms (I and II), as well as free GFP and GFP-mCherry forms. These forms indicate the autophagy flux. **(F)** In dopaminergic neurons, AUTEN treatments increased autophagy (accumulation of degraded GFP and GFP-mCherry products). On the box-plot diagrams, he black line indicates the median value, the box shows the most representative 50%, while filed black circles show the value of individual samples. Starts indicate P values where *: p < 0.05, ***: p < 0.005. Detailed description of the statistical methods used can be found in the Materials and Methods chapter.

When *Appl-Gal4* driver was used, only AUTEN-99 could significantly increase the amount of mCherry-Atg8a-positive structures in neurons of adult brain (Figure S3B). This result is highly similar to that obtained by using *Elav-Gal4*. The amount of GFP-2xFYVE-positive structures was significantly elevated when animals were treated with either of the AUTENs (Figure S3C). In dopaminergic neurons (*ple-Gal4*) and GABAergic neurons (*Gad1-Gal4*) neurons, the two AUTEN molecules also enhanced autophagic activity (the quantities of mCherry-Atg8a- and GFP-2xFYVE-labeled structures were elevated) (Figure 3A to 3D). In GABAergic neurons, an increase in the amount of GFP and GFP-mCherry proteins was visible which is indicative for an enhanced autophagic degradation (Figure 3F and Figure S4D). We found differences in the response to AUTEN treatments between cholinergic and glutamatergic/motoneurons. In cholinergic neurons, only AUTEN-67 was able to increase the amount of autophagic structures and degradation products (Figure 4 A, 4C and 4E), whereas in glutamatergic neurons and motoneurons, only AUTEN-99 proved to be effective (Figure 4B, 4D and 4F). It is worth mentioning that in case of cholinergic neurons, a coupled Western blot analysis indicated two types of degradation products (free GFP and GFP-mCherry), while then in glutamatergic neurons and motoneurons, only mCherry-GFP was detectable (Figure 4E, F and Figure S4E, F). It is possible that cholinergic and glutamatergic neurons possess different sets of acidic hydrolases. Based on this hypothesis, degradation of the GFP-mCherry protein into GFP and mCherry products may be more effective.

**Figure 4:**
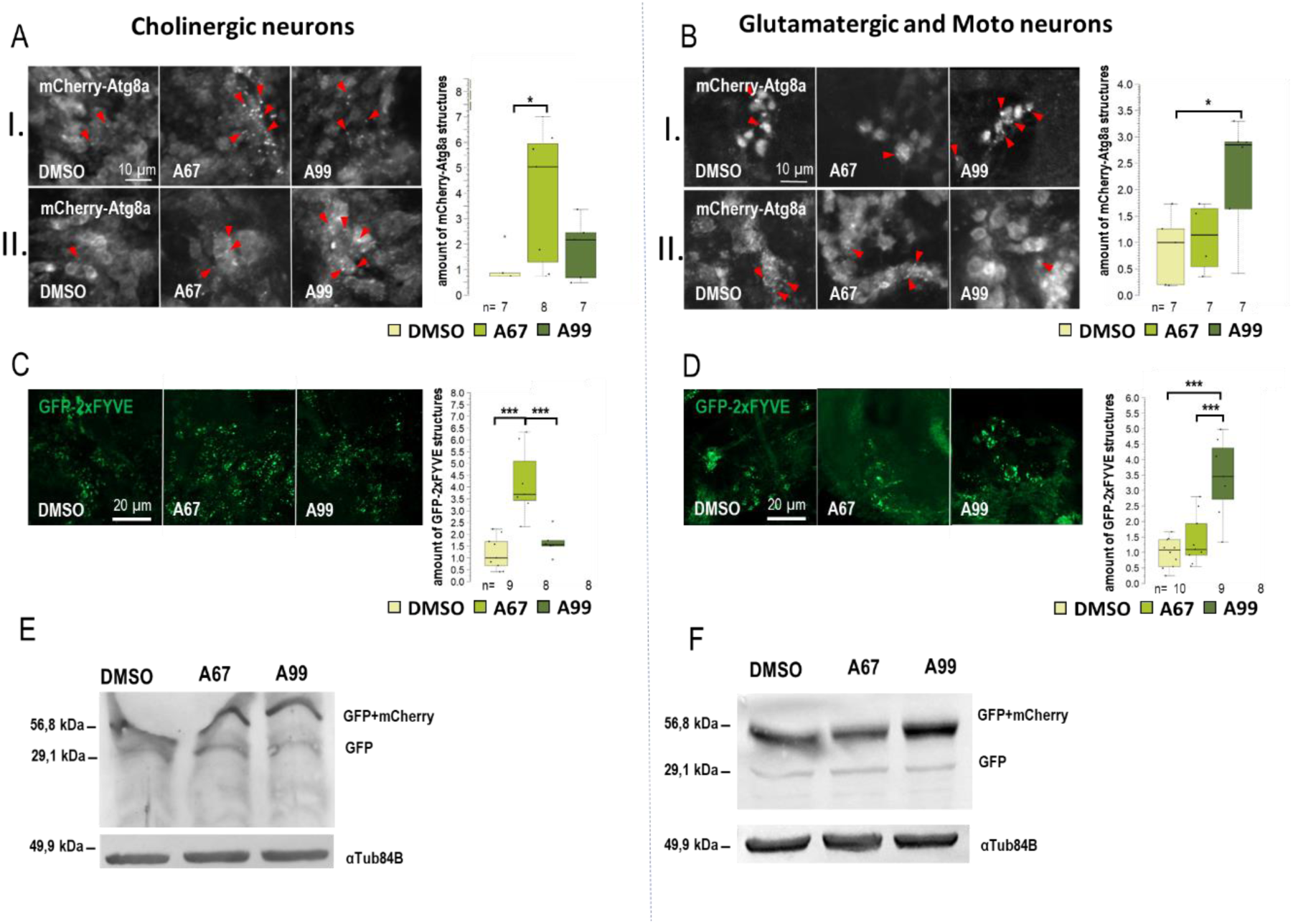
AUTEN-67 and AUTEN-99 regulate autophagy to different extents in cholinergic and motor neurons. The effect of AUTEN-67 (A67) and AUTEN-99 (A99) was monitored in a neuron-specific way. We examined the amount of autophagic vesicles in the brains of adult animals in two areas: (I) the mushroom body and (II) the neurons around the pharynx. **(A, B)** Expression of UAS-GFP-mCherry-Atg8a (UGMA) (red foci) in cholinergic (*Chat-Gal4* driver) and glutaminergic neurons/motoneurons (*OK371-Gal4* driver). (**A)** mCherry-Atg8a-positive structures (red) were identified by confocal microscopy. Red arrows indicate autophagic structures in cholinergic neurons. (**B)** The same labeling is observable in glutaminergic neurons and motoneurons. **(A-B)** In cholinergic neurons, A67 was able to significantly increase the amount of mCherry-Atg8a, while in glutamatergic/motor neurons, A99 did so. (**C-D)** Visualization of GFP-2xFYVE structures correlating with PIP3 levels. (**C)** A67 increases the amount of GFP-2xFYVE structures in cholinergic neurons. (**D)** A99 enhances PIP3 quantities in glutaminergic neurons and motoneurons only. (**E-F)** Expressing UGMA marker in a neuron-specific manner. The marker was used to quantify a GFP-mCherry-Atg8a reporter in cholinergic neurons (**G**) and glutaminergic neurons/motoneurons (**H**). Anti-GFP antibody was used for Western blotting. The ratio of GFP and GFP-mCherry proteins indicates the autophagy flux; the higher the ratio is, the more intense the process is. **(E)** Increased levels of free GFP proteins in cholinergic neurons treated with A67. **(F)** In glutamineric and moto neurons, A99 increased free GFP-mCherry levels. On the box-plot diagrams, the black line shows the median value, the box indicates the most representative 50%, while filled black circles display the value of individual samples. Stars indicate P values, where *: p < 0.05, ***: p < 0.005. Detailed description of the statistical methods used can be found in the Materials and Methods chapter.

### Both AUTEN molecules improve climbing ability and extends life span in GABAergic and glutamatergic neuron-specific SCA1 models

Next, we investigated the effect of the two AUTEN molecules on autophagy and physiology in flies accumulating mutant (82Q) ATXN1 proteins specifically in distinct types of neurons, including GABAergic, glutamatergic, moto- and cholinergic neurons (Figure 5, and Figure S5). 82Q ATXN1 significantly lowered moving ability in animals when it was expressed in any of these neuron types. In the GABAergic-specific SCA1 model, AUTEN-99 improved climbing ability in young adults only as compared with untreated control (Figure 5A). In the same model, AUTEN-67 positively influenced the ability of animals to climb at advanced ages only (Figure 5A). In the cholinergic-specific SCA1 model, AUTEN 67, but not AUTEN-99, increased locomotion (Figure 5B and Figure. S5B). These results are consistent with those obtained when studying the effect of the two AUTEN molecules on autophagy in specific neurons. Only AUTEN-67 could counteract defects in moving caused by overexpressing human mutant ATXN1 specifically in cholinergic neurons. DMSO-treated animals expressing ATXN1 exclusively in glutamatergic neurons and motoneurons died prior to age of 21 days (before the 3th test day). At the adult age of 14 days, in glutamatergic- and motoneuron-specific SCA1 models both AUTEN-67 and -99 could improve the ability of animals to climb up on the wall of a glass vial, as compared with untreated (only DMSO) control (Figure 5A and Figure S5).

**Figure 5.**
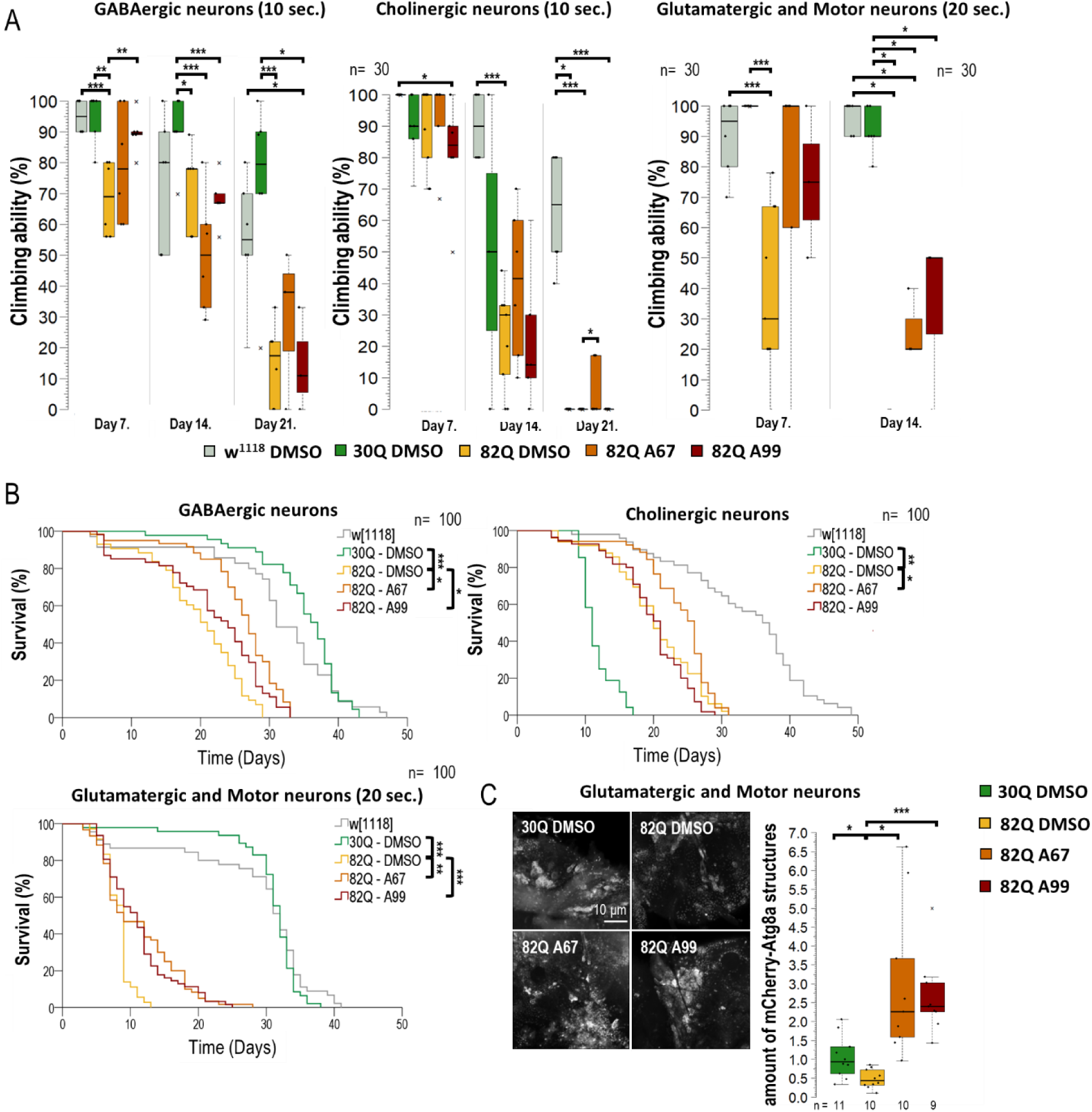
AUTENs improve climbing ability and extend life span in GABAergic, glutaminergic neuron-specific SCA1 models. The effect of AUTEN-67 (A67) and AUTEN-99 (A99) on the ability of animals to climb up on the wall of a glass tube. Animals were maintained at 29°C, and tests were performed at ages of 7, 14 and 21 days. Because 82Q animals were unable to climb up a 21.8 cm long distance within 1 minute, a 6 cm long distance was used for testing. The positions of animals were scored after 10 and 20 seconds. 82Q ATXN1 protein was expressed specifically in GABAergic, cholinergic or glutaminergic neurons. **(C)** 82Q ATXN1 expression in glutaminergic neurons and motoneurons shortened life span, so climbing assays in these cases were performed on animals with age of 7 and 14 days, and maintained at 29°C. 82Q control animals (only DMSO treated) died before age 14 days. Some of the 82Q animals treated with either of the AUTEN molecules lived for 14 days and retained its ability to move. **(B)** Surviving curves of neuron-specific SCA models. Expressing 82Q ATXN1 significantly interfered with survival in each of the cases. Both AUTEN molecules extended the life span of SCA1 models expressing 82Q ATXN1 in GABAergic or glutaminergic neurons. In the cholinergic neuron-specific CSA1 model, only A67 could promote longevity. Further relevant data are seen in Figures S5 and S6. **(C)** In glutamatergic neurons and motoneurons, A67 also reducesed the phenotype characteristic of SCA1 models. Therefore, we examined the specific expression of mCherry-Atg8a in glutamatergic and motor neurons treated with AUTEN in 82Q ATXN1 mutants, where both small molecules increased the number of autophagic vesicles **(C’)**. Our results suggest that both A67 and A99 can induce autophagy in glutamatergic-specific SCA1 models. On the box-plot diagrams, the black line represents median, while black-filled circles indicate the value of individual samples (at climbing, the percentage of 10 individuals).

A coupled life span assay provided results that are consistent with those obtained by the climbing test (Figure 5B and Figure S5). 82Q animals expressing mutant ATXN1 in GABAergic neurons only lived significantly shorter than 30Q and *w^1118^* control strains (Figure 5B and Fig. ure S5). Although both AUTEN molecules increased survival in these 82Q flies, AUTEN-67 appeared to be more effective in this regard. Expressing 82Q (mutant) ATXN1 proteins specifically in cholinergic neurons per se shortened life span, although 30Q ATNX1 expression in these neurons was also detrimental for survival (Figure 5B). In this model, AUTEN-67 was effective only (Figure 5B). In glutamatergic neuron- and motoneuron-specific SCA1 models, however, both AUTEN molecules promoted longevity (Figure 5B and Figure S5). In a previous measurement, we demonstrated that only AUTEN-99 can increase autophagy in glutamatergic neurons (Figure 3B, D, and F). However, in glutamatergic SCA1 models, both drugs improved survival and climbing ability. Therefore, we compared the effects of the two AUTEN compounds on autophagy in mutants overexpressing glutamatergic-specific 82Q ATXN1. Both AUTEN-67 and AUTEN-99 significantly increased the number of mCherry-Atg8a-positive vesicles (Figure 5C).

To sum, the two autophagy enhancers, AUTEN-67 and -99, positively influenced climbing ability and extended life span in GABAergic and glutamatergic neuron/moto neuron-specific SCA1 models despite the fact that AUTEN-67 appeared to be ineffective in activating autophagy in the latter model. In the cholinergic neuron-specific SCA1 model, however, only AUTEN-67 was effective.

### AUTENs modulate autophagy in mouse hippocampal neurons in a differential manner

We further investigated whether a differential effect of the two autophagy-activating small molecules could also be observed in a mammalian system. For this purpose, we utilized mouse hippocampal cell cultures derived from embryos aged 17-18 days. The cultures were treated with AUTEN-67 or -99 for a duration of 24 hours. To assess autophagy, we employed anti-LAMP1 and anti-p62 labeling (see Figure 6’). LAMP1 is a protein associated with the lysosomal membrane and serves as a marker for lysosomal vesicles, including primary lysosomes, autolysosomes, and endolysosomes [44]. Our findings demonstrated a significant increase in LAMP1 levels in hippocampal neurons following treatment with both compounds (Figure 6A’). However, AUTEN-99 resulted in a more pronounced decrease in p62 levels (Figure 6A’’). Notably, p62 levels were significantly reduced exclusively by AUTEN-99. As a negative control, an autophagy inhibitor – Bafilomycin – treatment is applied. Bafilomycin inhibits autophagosome-lysosome fusion as well as lysosomal acidification. During inhibition, we expect accumulation of p62 staining, thus assuming an opposite effect compared to AUTEN treatments. In Bafilomycin treated samples, a significant accumulation of p62 is observable (Figure 6B). From these results, we suggest that there may exist a distinction in the neuronal specificity of these drugs within mammalian cells. The embryonic hippocampal neuron cultures contained GABAergic neurons [45], which were by glutamic acid decarboxylase 65/67 (GAD65/67) immunostaining (Figure 6B). Interestingly, when p62 labeling was detected selectively in GABAergic cells (Figure 6B, B’), our results indicated that only AUTEN-99 treatment evoked a significant drop in p62 levels. This finding contradictssts with observations made in *Drosophila* GABAergic cells, where both drugs effectively increased the amounts of autophagic structures (Figure 3). However, our results confirm our previous more general observation that AUTEN small molecules can influence different neuron types to varying degrees.

**Figure 6.**
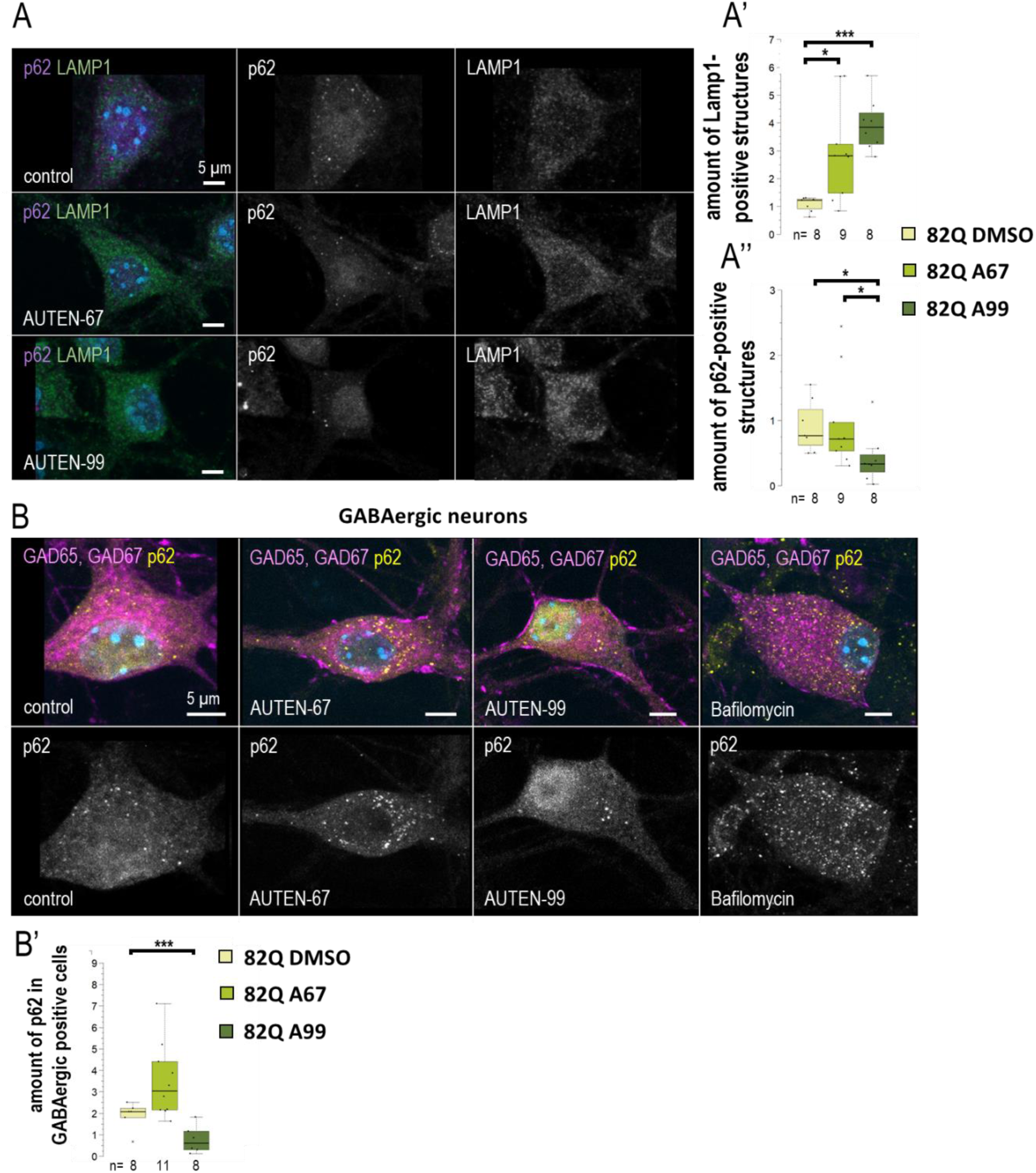
AUTEN-99 increases autophagy in GABAergic neurons. We investigated the effects of 24-hour AUTEN treatments on the number of lysosomes (LAMP1 – green) and the autophagy substrate (p62 – purple) in cultured mouse embryonic hippocampal neurons. Both compounds increased the amount of lysosomes, while p62 was significantly decreased only in the AUTEN-99 treated samples. **(B-B’)** Cultured hippocampal GABAergic neurons were labeled with anti-Gad 65 and -Gad 67 (magenta) and p62 (yellow). Only AUTEN99 reduced the amount of p62 structures in GABAergic cells. On the box-plot diagrams, the black line represents median, the box shows the most representative 50%, while black-filled circles indicate the value of individual samples (at climbing, the percentage of 10 individuals). For statistics, *: p < 0.05, **: p<0.01, ***: p < 0.005. Detailed description of the statistical methods used can be found in the Materials and Methods chapter.

## Discussion

When mutant (82Q) ATXN1 protein was expressed in the whole brain of a *Drosophila* SCA1 model, we detected significant differences in the ability of the two AUTEN small molecules to activate autophagy. AUTEN-67 treatment lowered the amount of ubiquitin- and p62/Ref(2)P-positive structures (both markers label autophagic substrates) and increased the level of 2xFYVE-GFP structures, thereby indicating an enhanced autophagic flux. Administration of AUTEN-99, however, elevated the number of mCherry-Atg8a-positive foci, whereas AUTEN-67 only tendentiously increased the amount of mCherry-Atg8a only tendentiously (Figure 1 and Figure S1).

Thus, AUTEN-99 may influence a larger brain area or trigger a more intense change in the accumulation of mCherry-Atg8a marker than AUTEN-67. Performing a Western blot analysis, both AUTEN molecules decreased p62/Ref(2)P levels, although AUTEN-67 did it at a much significant rate. AUTEN-67 and -99 also lowered the amounts of the lipid-conjugated Atg8a-II isoform more significantly as compared with soluble Atg8a-I. This indicates an elevated autophagic activity (Figure 1C) [33]. Despite their obvious functions in the brain, neuron-specific activity of these compounds remains unclear. The structure and size of the two AUTEN molecules highly differ from each other. Hence, it is possible that their capacity to penetrate through the cell membrane is cell type-specific.

We demonstrated previously that certain autophagy markers display a non-homogenous distribution in the brain and certain neuron types contain higher levels of autophagy structures than others [8]. Different types of neurons also vary in their capacity to take up, or to respond to, distinct active substances. Together, a combination of these factors is likely to underlie differences between the two compounds which could be detected when using the mCherry-Atg8a marker.

The two AUTEN molecules were able to enhance the life span of control animals expressing normal (30Q) ATXN1 proteins (Figure 2A and Fig. S2). Their effects were highly similar to each other in this genetic background. Contrary to these data, in animals expressing the mutant (82Q) ATXN1 protein only AUTEN-67 had the capacity to extend life span and improve climbing ability at advanced ages, whereas AUTEN-99 proved to be ineffective in this respect (Figure 2A, B and Figure S1A, B). Furthermore, AUTEN-67, but not -99, could reduce the number of abnormal ommatidia, which is a well-known characteristics of *Drosophila* SCA1 models [34] (Figure 2C and Figure S2C, C’) [34]. Thus, there are detectable differences in the effect of the two compounds to induce autophagy in SCA1 models, and this dissimilarity depends on the type of neurons in which the mutant ATXN1 protein is expressed. It is possible that AUTEN-67 and -99 penetrate into different neurons at distinct rates. Therefore, we also studied the effects of the two autophagy enhancers in a neuron type-specific manner (82Q ATXN1 was expressed in different types of neurons) (Figures 3 to 6). In patients diagnosed for SCA1, PCs of the cerebellum are mostly involved [46] which are regulated by other neurons through GABA and glutamate neurotransmitters [47]. Dendritic development in PCs also depends on GABAergic and glutaminergic innervations [47]. Although PCs represent GABAergic neurons, these cells control GABAergic interneurons [48]. The *Drosophila* brain does not contain PCs, but have GABAergic and glutamatergic neurons [49, 50]. The physiological role of the latter is still unclear. Cholinergic neurons in this organism are involved in visual perception [51]. Assaying these types of neurons is reasoned by the human SCA1 pathology [52]. Because in *Drosophila* SCA1 models, the mutant (82Q) ATXN1 protein has been expressed in the whole brain [27], here we specifically expressed the protein in different neuron types. Results we obtained show that both AUTEN molecules increased the autophagy flux in GABAergic (*Gad1-Gal4*) and dopaminergic (*ple-Gal4*) neurons (Figure 3). In mammalian cells, we also examined the effect of AUTENs on GABAergic neurons; however, AUTEN-99 was more effective in enhancing autophagy (Figure 6B, B’). This difference may be due to the fact that, in cell cultures, cells are directly influenced by the drug and do not need to pass through other tissues or the blood-brain barrier. AUTEN-67 and -99 could improve climbing ability in animals expressing mutant (82Q) ATXN1 protein. In contrast, AUTEN-67, but not AUTEN-99, was effective in animals expressing 82Q ATXN1 in cholinergic neurons only (*Chat-Gal4*) (Figure 4). When 82Q ATXN1 protein expression was restricted exclusively to glutamatergic neurons and motoneurons (*OK371-Gal4*), AUTEN-99, but not AUTEN-67, was able to exert a positive effect (Figure 4). However, both drug candidates cold increased life span in glutamatergic neuron-specific 82Q mutants (Figure 4). Further research may resolve this discrepancy. We previously found that Arl8 activation (through increased lysosome degradation) promotes survival in an otherwise wild-type genetic background, but did not influence life span in a Parkinson’ disease (PD) model (flies expressing human mutant alpha-synuclein, A53T) [53]. Differences between the effects of AUTEN-67 and -99 on climbing ability and life span may result from the fact that the latter cannot induce autophagy in neurons most exposed to SCA1 pathology. Alternatively, AUTEN-99 may hyperactivate the process in neurons that are most sensitive to autophagic degradation.

Alternatively, the rates at which the AUTEN molecules penetrate through the brain-blood barrier are dissimilar [22]. Further research should investigate whether these molecules also have unequal effects on different neurons in another genetic model. In addition, metabolites generated by the degradation of AUTEN-67 and -99 may also be different among distinct neurons. The observed differences in the effects of AUTENs raise further questions. The answers to these questions help us to decide when to use the two drug candidates.

In humans, defects in MTMR-14 function can lead to the incidence of centronuclear myopathy, which is a developmental muscle abnormality [54]. We earlier investigated the effects of the AUTEN compounds on striated muscle physiology in *Drosophila* [46]. The molecules improved flying ability and decreased the levels of protein aggregates and aberrant mitochondria, which are associated with muscle aging. Administrating these AUTENs led to no side effect, which was not true in case of EDTP deficiency [55]. Autophagy is regulated by factors acting downstream of vesicle nucleation. Generating new autophagosomes without promoting autolysosomal degradation could be harmful in senescent cells, in which the process has already been damaged at several stages [8]. MTMR14/EDTP proteins may also dephosphorylate phosphatydilinositol-3,5-bisphosphate (PI(3,5)P_2_) into phosphatidylinositol-5-sphosphate (PI5P), although this conversion step was previously linked to MTM, MTMR1 and MTMR6 proteins only [47, 48]. These lipid phosphatases are known to phosphorylate PI(3,5)P_2_ into PI5P [56, 57]. In *Drosophila*, EDTP hyperactivation in the nervous system increases the amount of the lipid-conjugated Atg8a-II isoform, indicating an additional function for this enzyme after Atg8a conjugation. This additional function could be the conversion of PI(3,5)P_2_ into PI5P [8]. In future, it would be worth investigating whether the AUTEN molecules have any effects downstream of vesicle nucleation.

The vesicle fusion process is regulated by HOPS (homotypic fusion and protein sorting), SNARE (soluble *N*-ethylmaleimide–sensitive factor attachment protein receptors) and small GTPase proteins [58]. Although the Rab2 (Ras-associated binding 2), Rab7 and Arl8 (ARF-like 8) small GTPases possess different membrane affinities, each of them is required for lysosomal breakdown [59–61]. We previously overexpressed constitutively active forms of small GTPases, and found that Rab2 and Arl8 hyperactivity enhance autophagy in wild-type and flies models of PD model flies [53]. Further investigation should address whether simultaneous activation of autophagy at multiple target sites can lead to a more effective effect. A parallel activation of vesicle fusion (MTMR14 inhibition) and nucleation (small GTPase activation) steps may cause a synergiestic effect.

Inhibiting EDTP/MTMR14 may also cause autophagy-independent effects. The Vps34 complex can link to UVRAG (UV radiation resistance-associated) protein. This molecular interaction controls the generation of PIP3-containing membranes during early endocytosis [62, 63]. According to results obtained from experiments on *Drosophila* larval fat body cells and HEK (Human Embryonic Kidney) cell cultures, neither EDTP nor the AUTEN molecules influence the endocytotic process [21–23]. The potential effects of EDTP and the AUTEN molecules on endocytosis have not been examined in neurons.

## Conclusions

We here suggest that both AUTEN-67 and -99 could be promising drug candidates in treating SCA1. These compounds may promote the survival and normal functioning of the affected neurons. The two AUTEN molecules have different effects on distinct neuron types. Further studies should clarify the molecular mechanisms underlying these differences.

## Materials and Methods

### *Drosophila* strains and AUTEN treatments

Animals were maintained under normal conditions on standard medium at 25°C. Treatments and experiments were carried out at 29°C. AUTEN-67 (ECHEMI Technology, T0501-7132) and AUTEN-99 (ECHEMI Technology, T0512-8758) were dissolved in DMSO (Sigma, D8418). Control animals were thus treated with DMSO only. For treatments, stock solutions were diluted to 100 μM in yeast suspension. Treated animals were placed on fresh drug medium every second day.

The following strains were obtained from the Bloomington *Drosophila* Stock Center (BDSC): *w[1118]* (BDRC 5905), *w[*]; P{w[+mC]=UAS-GFP-myc-2xFYVE}2* (BDRC 42712), *P{Appl-GAL4.G1a}1*, *y1 w\** (BDRC 32040), *w[*]; P{w[+mC]=ChAT-GAL4.7.4}19B* (BDRC 6793), *P{w[+mC]=Gad1-GAL4.3.098}2/CyO (BDRC 51630)*, *w[*]; P{w[+mC]=ple-GAL4.F}3* (BDRC 8848). The following strains were provided by the laboratory of Huda Zoghbi: *P{w[+mC]=UAS-Hsap\ATX1.30Q}F6 (30Q)*, *w[1118]; P{w[+mC]=UAS-Hsap\ATX1.82Q}M6 (82Q)*, *w[*]; P{w[+mC]=GAL4-elav.L}3*. From Gábor Juhász laboratory we got the next strains: *w[*]; P{UAS-mCherry-GFP-Atg8a}*, *w[1118]; 3xmCherry-Atg8a*, *w[*]; TubGFP-p62 3-2M (II)/CyO*, *w[*]; OK371-Gal4/CyO*.

### Immunohistochemistry

Experiments were performed according to the following protocol [53]. Brains isolated from females were fixed in 4% formaldehyde in PBST. For primary antibody labeling, samples were incubated in the primary antibody in a cold room for 2 days. Secondary antibody labeling was performed for 1 hour at room temperature. For nuclear labeling, Hoechst staining (dissolved in mounting medium) was used. Anti-ubiquitin (mouse, 1:500, Merck Life Science, ST1200). Anti-Rabbit Alexa Fluor 488 (Life Technologies, A11008) and anti-Mouse Texas Red (Life Technologies, T862) at 1:500 dilution were used as secondary antibodies.

### Fluorescence microscopy

For fluorescence microscopy, transgenic samples were fixed in 4% formaldehyde (diluted in PBS) for 20 min at room temperature. Mounting medium containing Hoechst was used to cover samples after washing the fixative solution (3×10 min with PBS). A Zeiss Axioimager M2 fluorescence microscope equipped with ApoTome was used for imaging (Department of Genetics at the ELTE University). For confocal imaging, a Zeiss inverted LSM800 confocal microscope was used (Institute of Biology at the ELTE University).

### Primary mouse hippocampal cultures

Wild-type CD1 mice were housed in the animal facility at 22 ± 1°C with 12-h light/dark cycles and ad libitum access to food and water. All experiments were carried out in compliance with local standards and laws regarding the use of experimental animals, as well as Hungarian and European Union legislation. Embryonic hippocampal cultures were prepared from CD1 mice on embryonic day 17-18 as described essentially by Oueslati Morales et al [45]. Cells were seeded onto poly-l-lysine -laminin–coated glass coverslips in 24-well plates in NeuroBasal PLUS (ThermoFisher Scientific; #A35829-01) culture medium supplemented by 2% B27 PLUS (ThermoFisher Scientific; #A3582801), 5% FBS (PAN Biotech; #P30-3309), 0.5 mM GlutaMAX (ThermoFisher Scientific; #35050-038), 40 μg/ml gentamicin (Sigma; #G1397), and 2.5 μg/ml amphotericin B (Life Technologies; #15290-026). Cultures were maintained in vitro for 11-12 days in 5% CO_2_ at 37°C prior to experiments, with changing one third of the medium to BrainPhys (StemCell Technologies; #05790) supplemented with 2% SM1 (StemCell Technologies; #05711), 40 μg/ml gentamicin, and 2.5 μg/ml amphotericin B medium on the 5th and 9^th^ days. Treatment with AUTENs (ECHEMI Technology, T0501-7132; ECHEMI Technology, T0512-8758) was applied on DIV 11 or 12, in a final concentration of 10 μM. In certain cases, cultures also received 50 nM Bafilomycin A1 (Merck; #B1793). 24 hours later, cultures were fixed with 4% paraformaldehyde in PBS for 20 minutes.

### Immunostaining and quantitative microscopy in fixed hippocampal cultures

Cultures were immunostained essentially as described by Bencsik et al. [64]Primary antibodies were anti-LAMP1 (mouse, 1:100; #1D4B; DSHB), anti-GAD65 (mouse, 1:6000, #GAD-6, DSHB), anti-GAD67 (mouse, 1:6000, # MAB5406, Merck), and anti-p62/SQSTM1 (rabbit, 1:2000, #P0067, Merck). Appropriate secondary antibodies were anti-mouse Atto 550 (1:500; #A21237, Merck), and anti-rabbit Alexa Fluor 633 (1:500; #A21070, Merck). Images were taken with an LSM 800 (Carl Zeiss) microscope with Plan Apochromat 63×/1.4 immersion objective.

### Western blot and protein isolation

Protein samples were obtained from the head of 21 days old adult females. Western blots were prepared according to the following protocol [53], and each measurement was repeated at least two times. Primary antibodies: anti-Ref(2)P/SQSTM1/p62 (rabbit, 1:2000 [65], anti-Atg8a (rabbit, 1:2000 [66], anti-ataxin (rabbit, 1:1000, Merck Life Science, C116639), anti-GFP (rat, 1:2500, Developmental Studies Hybridoma Bank, DSHB-GFP-1D2). Anti-Tub84B (mouse, 1:1000, Merck Life Science, T6199) was used as internal control. The following secondary antibodies were used: anti-rabbit IgG alkaline phosphatase (1:1000, Merck Life Science, A3687), anti-mouse IgG alkaline phosphatase (1:1000, Merck Life Science, A8438), anti-rabbit IgG alkaline phosphatase (1:1000, Merck Life Science, A8438). NBT-BCIP solution (Merck Life Science, 72091) diluted in 3% milk powder TBST was used to develop antibody labeling. We used ImageJ software to analyze the western blot bands. We simultaneously determined the changes in the levels of free (degraded) mCherry-GFP and GFP. Their quantities were corrected for variations in the anti-Tub84B signals. We repeated the western blots at least three times.

### Life span measurements

For life span measurements, control and treated animals were placed on a new active medium every other day. The number of dead males and females was counted every day. 5 parallel tubes of each species were measured, with each tube containing 15 males and 15 females. SPSS 17 software was used for statistical evaluation.

### Climbing ability test

For climbing assays, animals were placed in a long, thin glass tube (25 cm high, 1.5 cm in diameter) using CO_2_ anesthesia. A recovery time of 1.5 h before the measurement was allowed. Negative geotaxis was induced by gentle tapping, which caused the animals to fall down the bottom of the tube, and they then move upwards. The number of animals reaching the line at 21.8 cm within 20, 40 and 60 seconds was measured. Two parallel tubes (15-15 animals) were used for each treatment, and measurements were repeated three times with half-hour breaks.

### Quantification and statistical analysis

All datasets were first tested for normality, using Shapiro-Wilk test. If they were, F-test was applied to verify variance, and if the variance was also equal, data sets were compared by using two-sample t-test. If the variances did not match, Welch-test was used. When one or both compared data sets were not normally distributed, Man-Whitney U-test was performed to compare samples. RStudio version 2023.06.2 was used for the analysis.

## Ethics approval

This study was conducted under the approval of the Institutional Animal Ethics Committee of Eötvös Loránd University (approval number: PEI/001/1108-4/2013 and PEI/001/1109-4/2013). All methods were performed in accordance to the guidelines on research ethics of Eötvös Loránd for the use of experimental animals, in agreement with European Union and Hungarian legislation

## Availability of data and materials

The datasets used and/or analysed during the current study are available from the corresponding author on reasonable request.

## Author Contributions

T.B. and M.A. performed experiments on *Drosophila* samples, analysed data and wrote the manuscript. F.K. and T.S. performed experiments on *Drosophila* samples and analysed data. N.B. and K.S. carried out experiments on primary hippocampal neurons and analysed data. V.A.B. designed experiments. T.V. and T.K. designed experiments, analyzed data, and wrote the manuscript.

## Acknowledgments

*Drosophila* strains and reagents were kindly provided by Gábor Juhász (HUN-REN Biological Research Centre Szeged, Szeged, Hungary) and Huda Zoghbi (Baylor College of Medicine, Houston, TX 77030, USA). The authors also thank Regina Preisinger, Beatrix Supauer, Erzsébet Gatyás for the excellent technical assistance.

